## Supplementary tables and figures for "HAP40 orchestrates huntingtin structure for differential interaction with polyglutamine expanded exon 1"

### Supplementary Information

**Supplementary Table 1. Cryo-EM data collection, refinement and validation statistics**

| | HTT-HAP40 (EMDB-22106) (PDB 6x9o) |
|---|---|
| **Data collection and processing** | |
| Magnification | 165,000 |
| Voltage (kV) | 300 |
| Electron exposure (e–/Å$^2$) | 48.0 |
| Defocus range (μm) | 1.2 – 3.0 |
| Pixel size (Å) | 0.822 |
| Symmetry imposed | C1 |
| Initial particle images (no.) | 2,240,373 |
| Final particle images (no.) | 647,468 |
| Map resolution (Å) | 2.6 |
| FSC threshold | 0.143 |
| Map resolution range (Å) | 2.5-3.5 |
| **Refinement** | |
| Initial model used (PDB code) | 6EZ8 |
| Model resolution (Å) | 2.6 |
| FSC threshold | 0.143 |
| Model resolution range (Å) | 2.5-3.5 |
| Map sharpening *B* factor (Å$^2$) | -42.3 |
| Model composition | |
| Non-hydrogen atoms | 20899 |
| Protein residues | 2669 |
| Ligands | 0 |
| *B* factors (Å$^2$) | |
| Protein | 47.14 |
| Ligand | N/A |
| R.m.s. deviations | |
| Bond lengths (Å) | 0.006 |
| Bond angles (°) | 1.102 |
| Validation | |
| MolProbity score | 2.11 |
| Clashscore | 14.85 |
| Poor rotamers (%) | 0.56 |
| Ramachandran plot | |
| Favored (%) | 93.41 |
| Allowed (%) | 6.29 |
| Disallowed (%) | 0.30 |

**Supplementary Table 2. Residues contributing to putative ligand-able pocket at interface of HTT and HAP40**

| | Pocket Contributing Residues |
|---|---|
| HTT | L1015, R1021, T1024, M1071, T1074, L1075, S1078, W1080 |
| HAP40 | P84, A87, L88, T91, E92, R95, H132, Q137, A139, A140, A143, L144, L146, E147, A150, R153, F165, E186 |

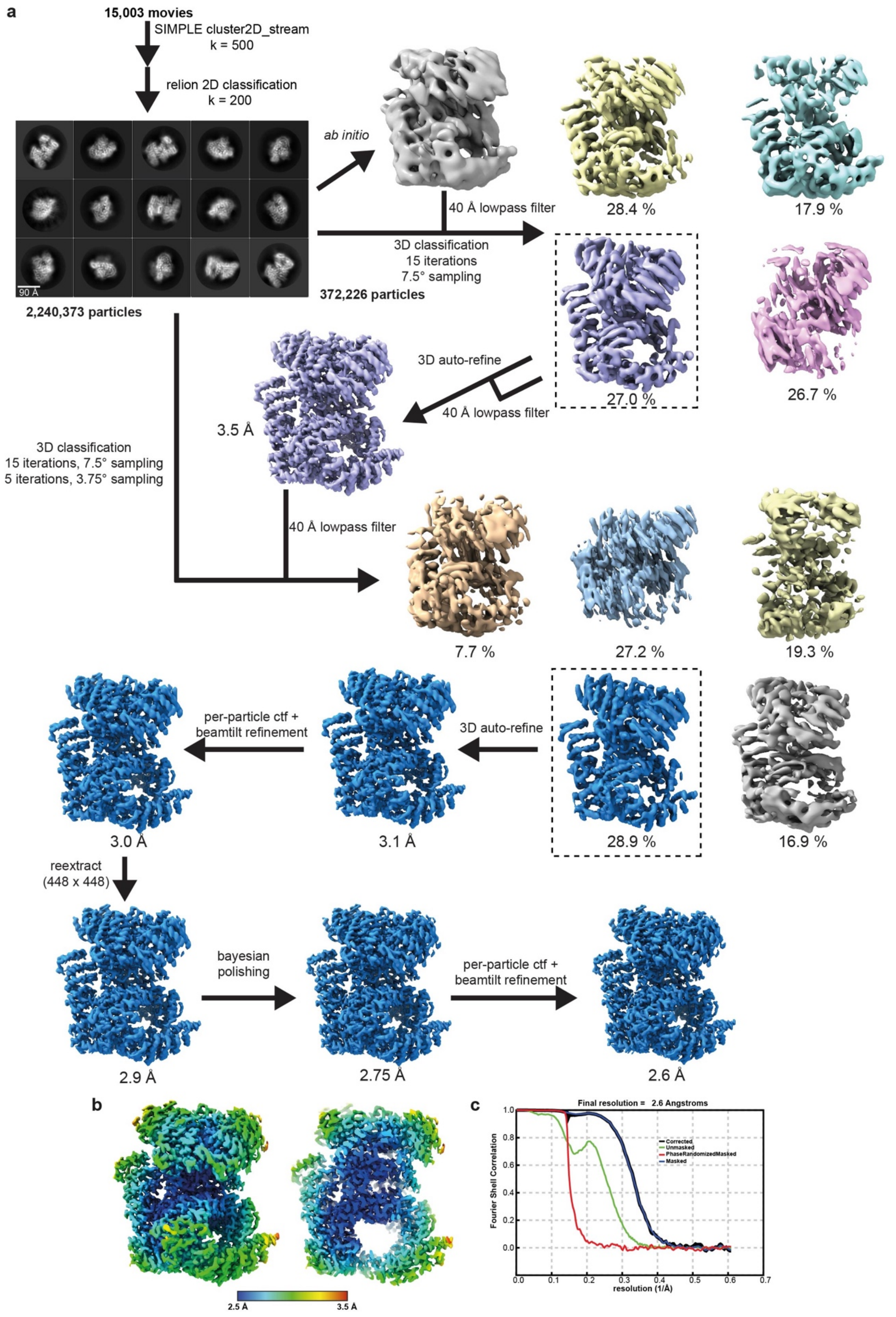

**Supplementary Figure 1. Cryo-EM processing workflow and local/global map quality for HTT-HAP40.** **a** Image processing workflow for HTT-HAP40. **b** Local resolution estimation of reconstructed map as determined within RELION. Left, full map; right, central slab through map. **c** Gold-standard Fourier Shell Correlation (FSC) plot used for global resolution estimation as determined within RELION.

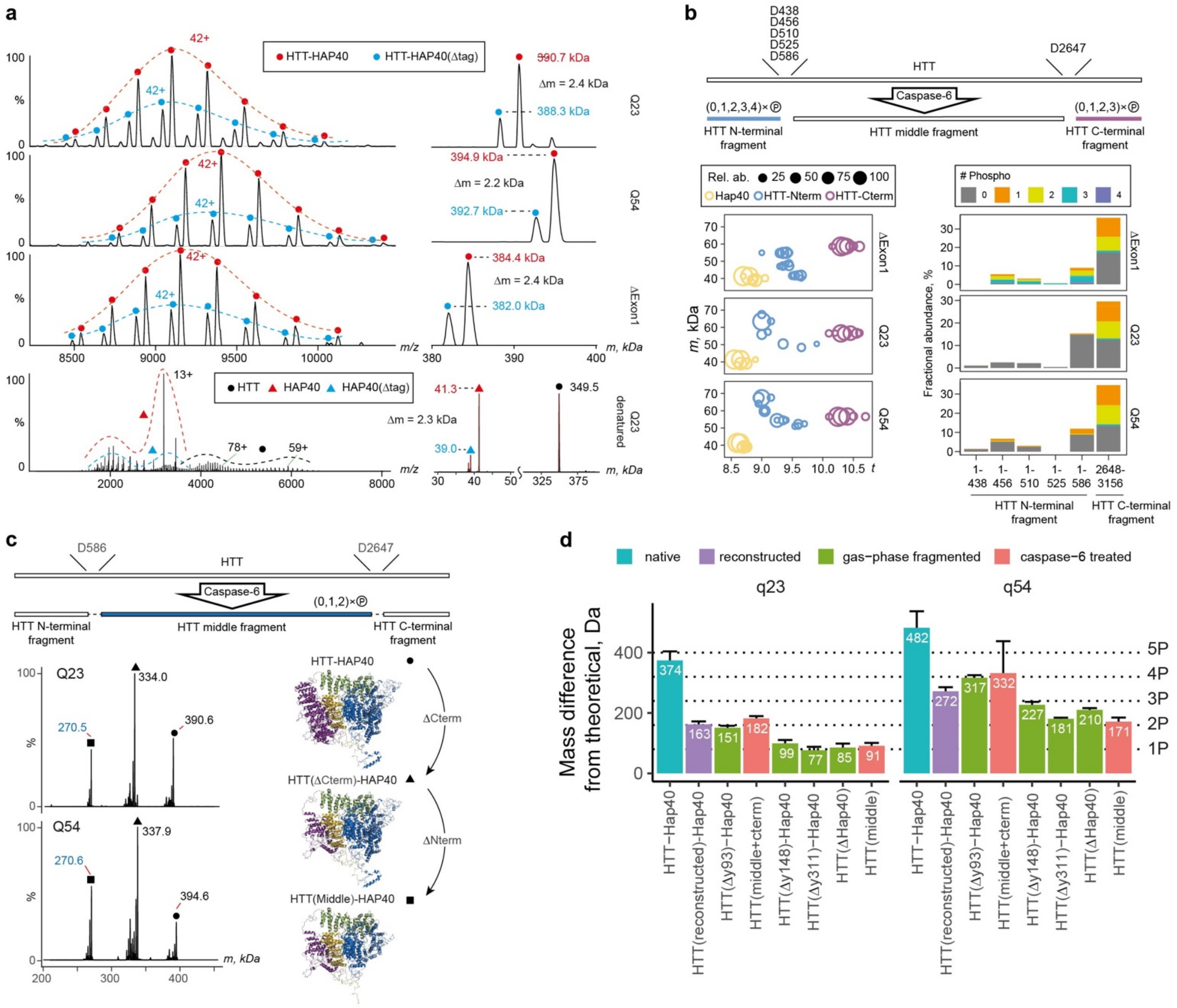

**Supplementary Figure 2. Determining the different proteoforms of HTT using hybrid mass spectrometry approach**.

**a** Native mass spectra for HTT-HAP40 Q23, Q54 and Δexon 1 samples. Top three left panels: annotated native raw spectra of HTT-HAP40 Q23, Q54 and Δexon 1. Top three right panels: annotated mass distributions of HTT-HAP40 Q23, Q54 and Δexon 1. Bottom panel: denaturing MS of HTT-HAP40 Q23 and respective mass distribution. Denaturing MS assigns the second peak, which is ~2.4 kDa smaller than the main peak and corresponds to the complex with the N-terminal His-tag cleaved from HAP40. **b** Middle-down MS of Caspase6-treated HTT-HAP40 complexes. Top panel: schematics of HTT digestion using Caspase6. Bottom left panel: mass-feature maps for digested and denatured HTT-HAP40 samples. Bottom right panel: abundances of N-terminal and C-terminal HTT proteoforms with 0-4 phosphorylation motifs (assigned by mass). **c** Native MS of Caspase6-treated HTT-HAP40 Q23 and Q54 complexes. Top panel: schematics of HTT digestion using Caspase6 with the uncleaved HTT-middle region highlighted. Bottom left panel: mass distributions of Caspase6-digested and gas-phase-activated HTT-HAP40. Bottom right panel: structures of HTT-HAP40 for the major species observed in native spectra on the left. **d** Difference between the theoretical and experimental masses of HTT-HAP40 complex samples under various conditions (native, Caspase6-treated and denatured, and gas-phase fragmented). Reconstructed masses were obtained by summing masses of HTT-middle determined by native MS and intensity-weighted average masses of HTT C-terminal and HTT N-terminal fragments produced upon Caspase6 treatment and determined in middle-down MS. Error bars represent mean error of mass determination.

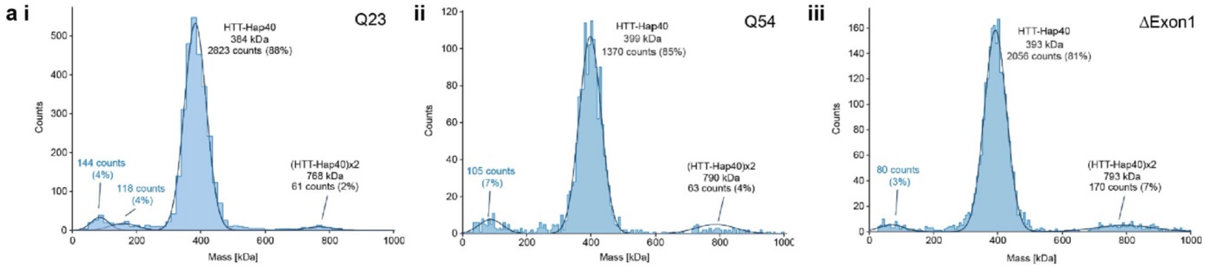

**b**

Starting fluorescence (AU)

| Buffer (all 50 mM) | pH | HTT-HAP40 Q23 NaCl (mM) 0 | 100 | 300 | 1000 | HTT-HAP40 Q54 NaCl (mM) 0 | 100 | 300 | 1000 | HTT-HAP40 ΔExon1 NaCl (mM) 0 | 100 | 300 | 1000 |
|---|---|---|---|---|---|---|---|---|---|---|---|---|---|
| Sodium Acetate | 4.8 | 92557 | 112657 | 38655 | 46969 | 109095 | 94575 | 24604 | 21194 | 63962 | 93927 | 23071 | 48598 |
| Bis Tris | 5.5 | 57918 | 23501 | 24669 | 54526 | 21774 | 14089 | 12908 | 27316 | 29922 | 21527 | 18050 | 24597 |
| Bis Tris Propane | 6 | 18487 | 13067 | 16939 | 14363 | 10479 | 9840 | 11142 | 7404 | 8334 | 8717 | 7365 | 11196 |
| HEPES | 7.4 | 11569 | 15886 | 15443 | 12805 | 12492 | 10260 | 8291 | 6804 | 7733 | 8527 | 6272 | 7651 |
| Tris | 8 | 19765 | 17057 | 11814 | 13731 | 10664 | 10155 | 8119 | 9598 | 7532 | 8260 | 7174 | 8747 |
| Tris | 8.5 | 15905 | 16355 | 7908 | 15257 | 7237 | 14516 | 5901 | 9236 | 9251 | 10037 | 6252 | 8235 |
| Bis Tris Propane | 9 | 18382 | 18455 | 17536 | 13984 | 10854 | 13187 | 13099 | 11758 | 9021 | 7824 | 10152 | 9322 |

Transition temperature (°C)

| Buffer (all 50 mM) | pH | HTT-HAP40 Q23 NaCl (mM) 0 | 100 | 300 | 1000 | HTT-HAP40 Q54 NaCl (mM) 0 | 100 | 300 | 1000 | HTT-HAP40 ΔExon1 NaCl (mM) 0 | 100 | 300 | 1000 |
|---|---|---|---|---|---|---|---|---|---|---|---|---|---|
| Sodium Acetate | 4.8 | <20.0 | <20.0 | <20.0 | <20.0 | <20.0 | <20.0 | <20.0 | <20.0 | <20.0 | <20.0 | <20.0 | <20.0 |
| Bis Tris | 5.5 | 37.2 | 36.3 | 34.6 | <20.0 | 34.6 | 37.7 | 36.6 | 33.6 | 38.2 | 36.4 | 35.3 | 32.9 |
| Bis Tris Propane | 6 | 41.8 | 42.3 | 41.9 | 41.3 | 43.5 | 43.8 | 43.0 | 42.9 | 42.5 | 43.3 | 43.2 | 42.7 |
| HEPES | 7.4 | 45.1 | 46.5 | 47.4 | 45.8 | 46.9 | 47.7 | 48.1 | 48.0 | 45.2 | 47.0 | 48.6 | 46.8 |
| Tris | 8 | 46.0 | 45.9 | 45.6 | 45.1 | 47.4 | 47.7 | 48.8 | 47.8 | 46.5 | 47.3 | 47.8 | 46.4 |
| Tris | 8.5 | 43.8 | 44.7 | 44.5 | 44.5 | 45.5 | 48.0 | 47.8 | 46.6 | 43.8 | 46.3 | 47.1 | 46.0 |
| Bis Tris Propane | 9 | 44.3 | 45.1 | 44.4 | 43.9 | 46.2 | 46.8 | 47.1 | 46.7 | 45.0 | 47.2 | 45.9 | 45.4 |

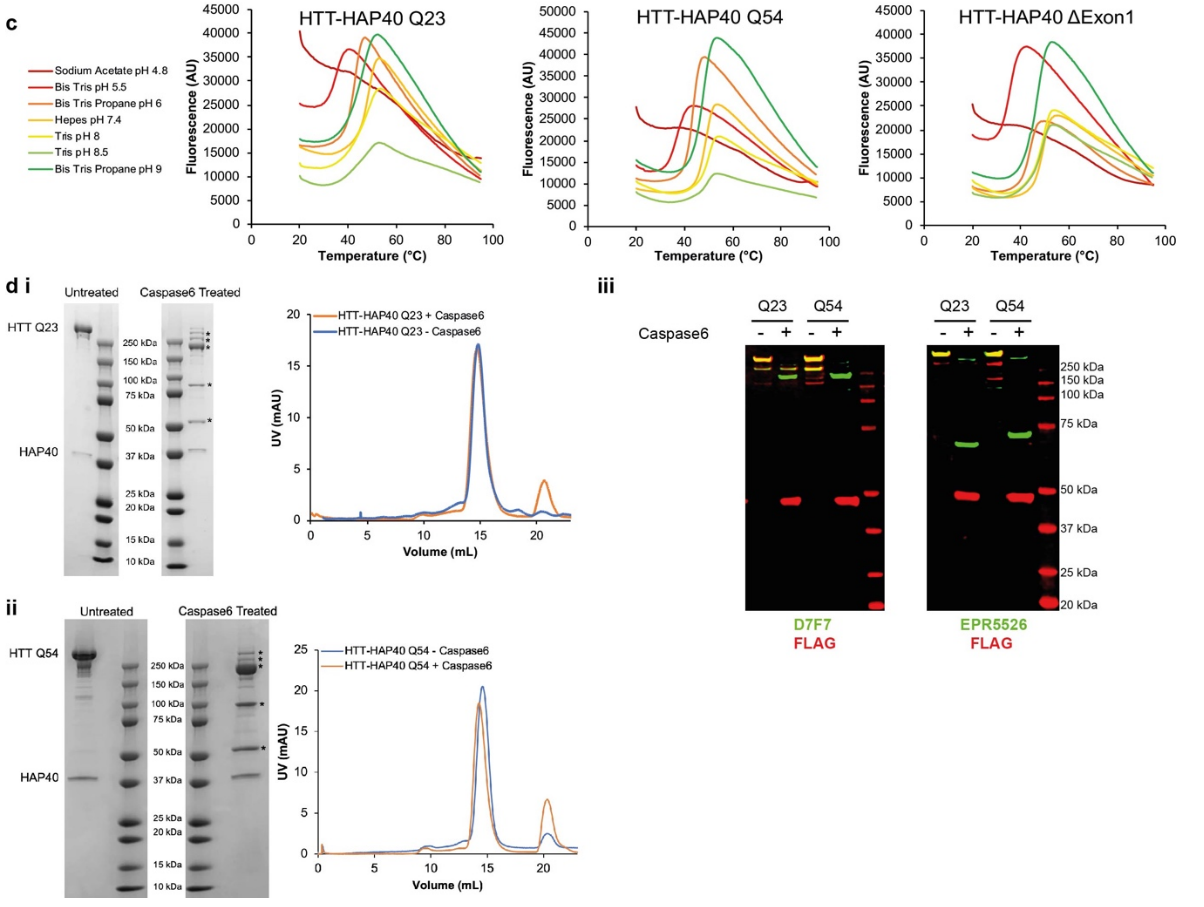

**Supplementary Figure 3. Stability and monodispersity of HTT-HAP40 complexes probed with buffer screening and proteolytic cleavage.**

**a** Mass photometry analysis of HTT-HAP40 Q23, Q54 and Δexon 1. **b** Assessing complex stability by measuring transition temperature using DSF in a range of different buffer and salt conditions. **c** DSF profiles of HTT-HAP40 samples in different buffer conditions with 300 mM NaCl. **d** Caspase6 cleavage of **i** Q23 and **ii** Q54 HTT-HAP40 proteins assessed by SDS-PAGE (left), analytical gel filtration (right) and **iii** western blot with antibodies recognising before (EPR5526) and after (D7F7) the documented Caspase6 cleavage site (D586) as well as anti-FLAG which recognises the C-terminal FLAG-tag of the samples.

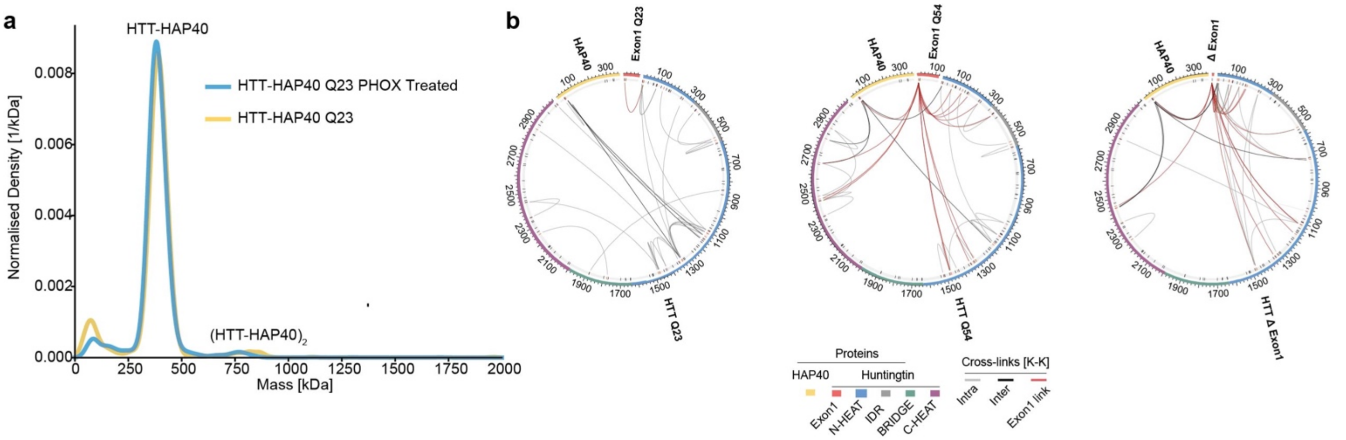

**Supplementary Figure 4. Mass photometry and cross-linking mass spectrometry analysis of HTT-HAP40.**

**a** Mass photometry analysis of HTTQ23-HAP40 untreated or treated with PhoX (0.5 mM). **b** Unique cross-links for each HTT-HAP40 sample with exon 1 in red, N-HEAT in blue, bridge domain in green, IDR in grey, C-HEAT in purple and HAP40 in yellow. Cross-linked lysine residues are indicated in red and unmodified lysine residues are indicated in black on the numbered sequence. Intermolecular cross-links (HTT-HAP40) are shown in black, intramolecular cross-links (HAP40-HAP40 or HTT-HTT) are shown in grey and exon 1 cross-links are shown in red. All residues following the exon 1 region of the different constructs are numbered the same for clarity.

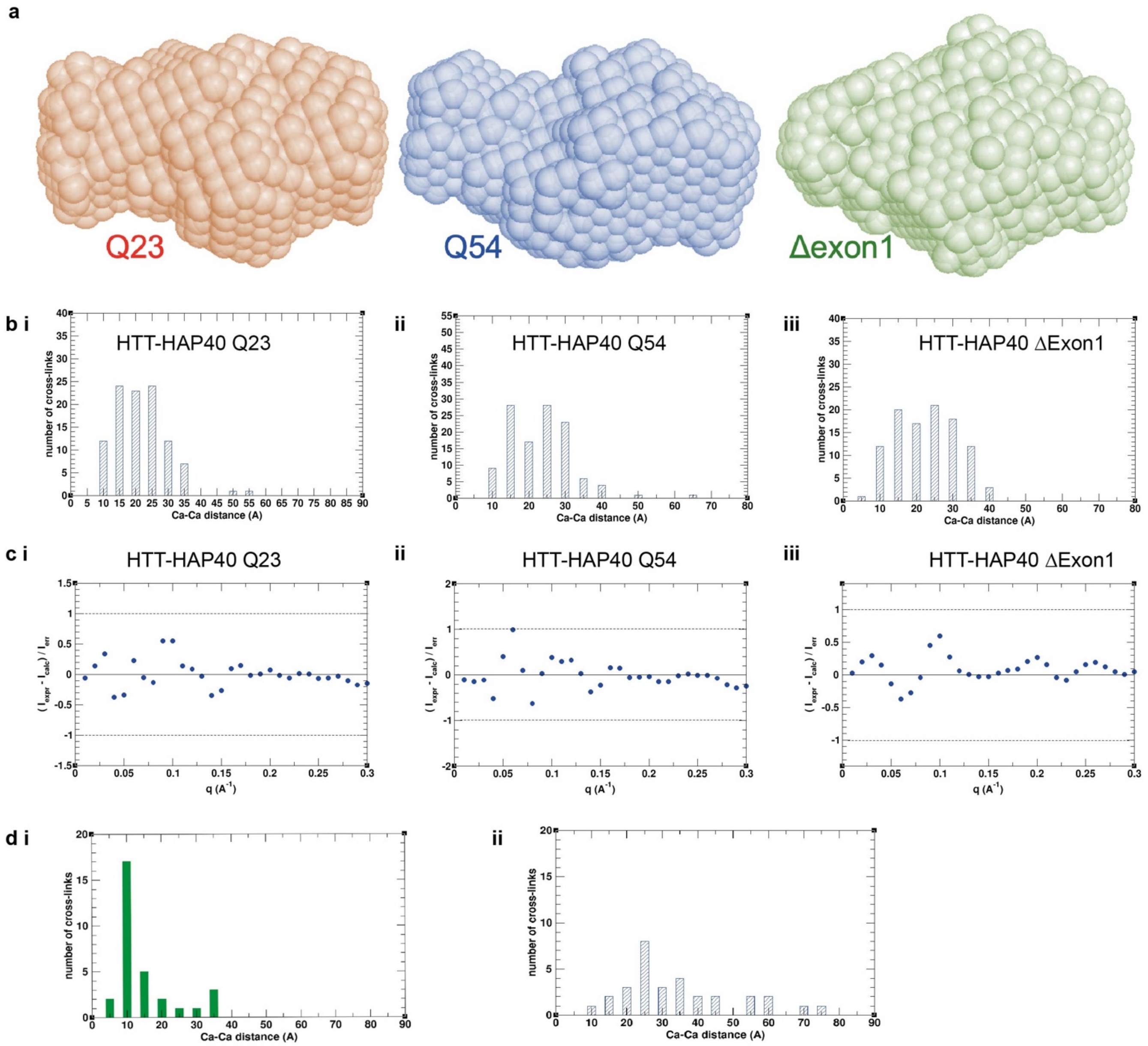

**Supplementary Figure 5. SAXS and cross-linking mass spectrometry analysis of HTT-HAP40.**
**a** SAXS envelopes calculated for HTT-HAP40 Q23 (red), Q54 (blue) and Δexon 1 (green). **b** Consistency of the experimental cross-links with an ensemble of models. Each histogram bar shows number of cross-links that have the corresponding Cα-Cα distances (the minimal distance over all models in the ensemble) falling within the corresponding distance bin. The histogram bars corresponding to the distances of > 35 Å indicate the number of cross-links that are inconsistent with the ensemble. **c** Normalized difference between the experimental SAXS profile and the theoretical profiles averaged over the ensemble. **d** Consistency of Q54 cross-links involving exon 1 region with two ensembles of Q23. The histogram bars for Cα-Cα distances > 35A indicate number of cross-links not consistent with an ensemble. **i** Q23 pool ensemble (18,000 models) is consistent with all cross-links; **ii** Q23 optimal ensemble (19 models) is inconsistent with 10 cross-links (including cross-links to C-HEAT domain). All cross-links are consistent with the pool, meaning that Q23 exon 1 region can form all cross-links that were observed for the Q54 exon 1 region.
